# Knowledge and interactions of the local community with the herpetofauna in the forest reserve of Quininí (Tibacuy, Cundinamarca)

**DOI:** 10.1101/814293

**Authors:** Juan Camilo Ríos-Orjuela, Nelson Falcón-Espitia, Alejandra Arias-Escobar, María José Espejo-Uribe, Carol Tatiana Chamorro-Vargas

**Affiliations:** Museu de Zoologia da Universidade de São Paulo, Avenida Nazaré 481, Ipiranga, São Paulo, SP CEP 04263-000, Brasil; Grupo de Morfología y Ecología Evolutiva, Instituto de Ciencias Naturales, Universidad Nacional de Colombia, Sede Bogotá, Apartado 7495, Bogotá D.C., Colombia; Laboratorio de Ecología Evolutiva, Departamento de Biología, Facultad de Ciencias, Universidad Nacional de Colombia, Sede Bogotá, Ciudad Universitaria, Bogotá D.C. 11001, Colombia; Grupo estudiantil de Herpetología, Área curricular de Biología, Facultad de Ciencias, Universidad Nacional de Colombia, Sede Bogotá, Ciudad Universitaria, Bogotá D.C. 11001, Colombia; Grupo de Biodiversidad y Conservación Genética, Instituto de Genética, Universidad Nacional de Colombia, Sede Bogotá, Ciudad Universitaria, Bogotá D.C. 11001, Colombia

**Keywords:** Ancestral knowledge, ethnoherpetology, herpetofauna, interactions, local communities

## Abstract

**Background:** The study of human-nature relationship has made possible to understand the life dynamics of the communities and the biodiversity with which cohabits. Although ethnobiological studies have been rise over the last decade, little is known about human interaction with herpetofauna in South America and in Colombia. In this work, we analyzed the knowledge, perception, and interaction of a local community located in the forest reserve of Quininí (RFPCQ) in Cundinamarca (Colombia), with respect to the herpetofauna that inhabits the area.

**Methods:** We performed semi-structured surveys containing 30 questions categorized into three groups: academic knowledge (1), use and cultural beliefs (2) and interaction and perception (3) related to the herpetofauna that occurs in the region. For the obtained data in question groups 1 and 2, an analysis and classification of the answers in percentages were made. For the question group 3, we assigned the answers with a hostility value according to the possible reaction of each individual interviewed in a hypothetical encounter with the herpetofauna, and performed a Multivariate Ordinal Logistic Regression test (MOLR), in order to know if the positive or negative reactions could be predicted by demographic variables.

**Results:** The community recognized the presence of amphibians and reptiles that cohabit their space, as well as their potential habitats. In addition, the role of herpetofauna was recognized in the magical/religious traditions for some inhabitants of the region, mainly associated with the fate and cure of chronic diseases. In general, the perception of amphibians and reptiles varied according to the origin and gender of the people, tend to have a more positive perception about reptiles than amphibians in most cases.

**Conclusions:** Although there was a general lack of knowledge on the part of the inhabitants of the RFPCQ about the biological and ecological aspects of herpetofauna, the population recognized the basic information about the habitats of these animals within the area of the reserve. There is a wide variety of uses of amphibians and reptiles in traditional medicine. Greater efforts should be made in the transmission and dissemination of knowledge about the ecological functions of herpetofauna.

## Background

Human communities have established a close relationship with the herpetofauna with which they cohabit, based on the use and understanding of amphibians and reptiles [1,2]. Thereby, the ethnoherpetology (as an integral part of ethnobiology) studies traditional knowledge between human-herpetofauna relationships, such as how a social group classifies and identifies different amphibians and reptiles species and their traditional uses [3].

The *Reserva Forestal Protectora Cerro Quininí* (RFPCQ) forest reserve is a conservation area located in Sumapaz province (Cundinamarca, Colombia), which is one of the most important biological corridors not only for the region but also for the eastern Andes, known as the Chingaza-Sumapaz-Guerrero corridor [4]. In addition, records of archaeological remains, such as petroglyphs and passage routes, associated with Panche community are known [5,6] which was an indigenous culture that inhabited this territory in pre-colonial times.

Due to its cultural and biological importance, RFPCQ has been focus of studies in diversity [7–12], ecology [13–15], anthropology [5,6,16–18], rural development and innovation in production systems [19,20], among others. Although most of the research associated with RFPCQ remains unpublished literature, these have actively involved community participation, which allows them to appropriate environmental aspects. This initial approaches to environmental problems constitute a fundamental basis for understanding the current ecosystems dynamics.

Since some years ago, ethnozoological studies have been made in Colombia, focused mostly on reptiles with high commercial usage, e.g. *Trachemys venusta callirostris*, *Iguana iguana*, *Boa constrictor*, *Tupinambis teguixin*, among others [21–24]. Nevertheless, there is poor information about the use and perception of herpetofauna, which concerns if the high biological diversity of Colombia and Andes mountains is taken into account [25–27]. In this way, efforts to obtain information about the fauna and its interactions within the habitat become relevant, where ethnobiological studies play a fundamental role in understanding the recent ecosystem dynamics, such as changes in the vegetal covertures and biodiversity among others [28,29].

Colombia is one of the most diverse countries of the world in herpetofauna species; currently, 616 species of reptiles and 835 species of amphibians [30,31] are registered in Colombia; with 115 and 367 endemic species, respectively [27]. Given this scenario, it is important to understand local communities’ perception about the species with which they share their territory. This may give us information about the historic relationship of the human groups with nature and can be the base for establishing conservation plans at local and regional scale that involve academy, politics and local communities [32].

Taking this into account, our aim is to evaluate the knowledge and perception of the RFPCQ community in relation with the herpetofauna in three different aspects: (1) academic knowledge, (2) use and cultural beliefs, and (3) human-herpetofauna interactions. In this way, we propose an initial approach that could clarify the relationship between the community and amphibians and reptile species that inhabit their territory.

## Methods

### Study area

The *Reserva Forestal Protectora Cerro Quininí* (RFPCQ) forest reserve is located in Tibacuy, Nilo and Viotá municipalities (Cundinamarca, Colombia) (Fig. 1), between 1050 and 2133 m above sea level on Eastern Andes range. In this region, the temperate climate predominates, with an average annual of 19.2 °C and a bimodal rainfall pattern [33]. With an approximate area of 1947 ha, it is one of the most extensive conservation areas in Cundinamarca department [4].

**Fig. 1.**
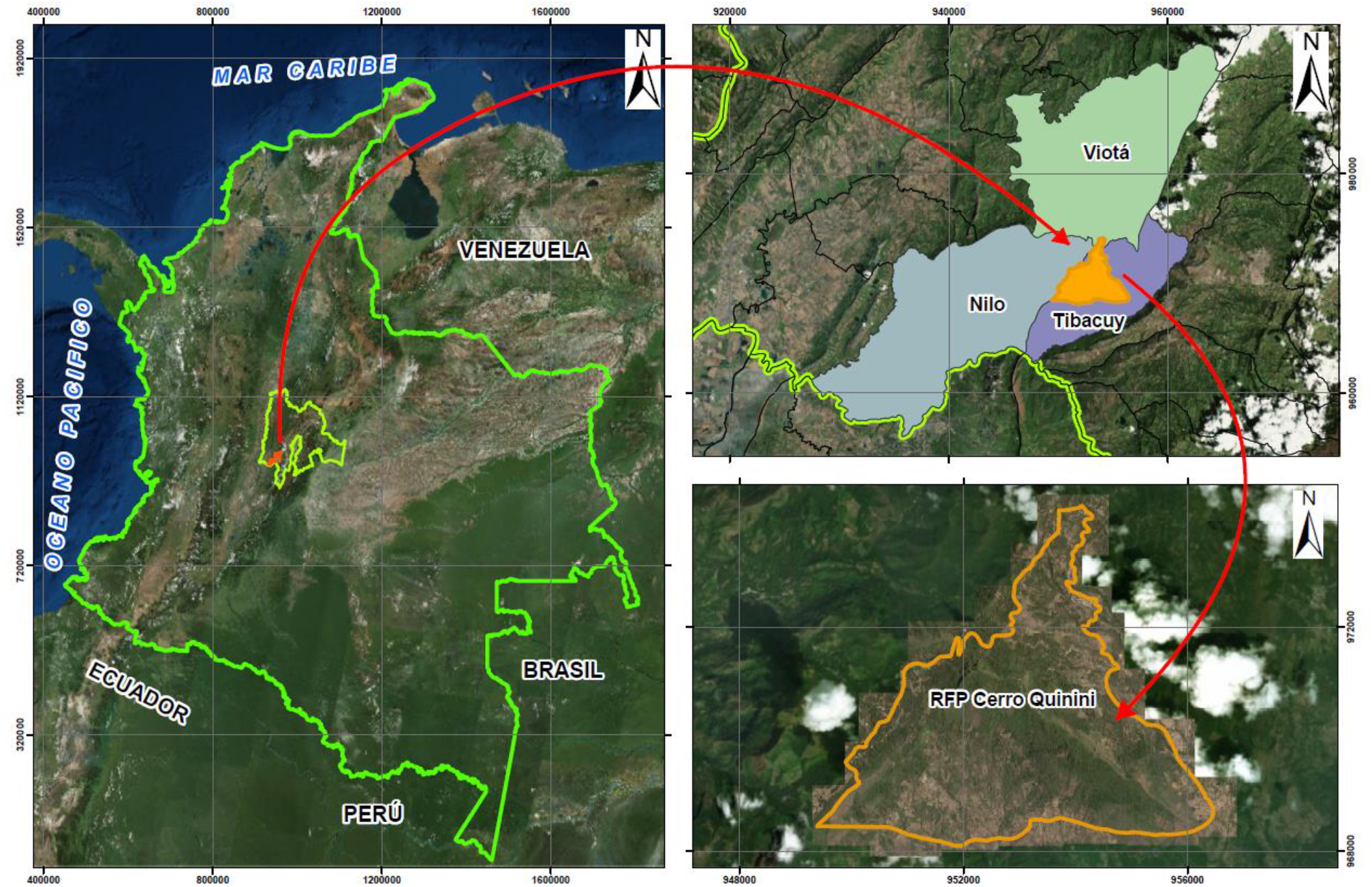
Localization of *Reserva Forestal Protectora Cerro Quininí* (RFPCQ) forest reserve.

Following the classification of Holdridge (1947) [34], the RFPCQ is located in the life zones of lower mountain moist forest and lower mountain dry forest to a lesser extent. Despite its protected area status, about 90% of the land associated with the RFPCQ are private properties, mainly with mixed crops, grazing areas and coffee crops being the largest [20,33,35,36].

This area was declared a protective forest reserve in 1987 as a measure for the conservation of the renewable natural resources and the environment by request of the people inhabiting the area, because of the important water births that supply the region [37].

### Data collection

Following Margoluis & Salafsky (1998) and Drumond et al. (2009) [38,39], we made 61 semi-structured surveys during April 2016 and June 2019 to randomly selected respondents on the communities of Cumaca inspection, Albania and La Vuelta villages (Tibacuy municipality), gathering information from people in urban and rural areas near to the RFPCQ.

Each survey had 30 questions of which 12 and 18 questions were related to general topics of amphibians and reptiles, respectively. We take demographic information like name, gender, age, occupation, birthplace, and education level in the following categories: illiterate, fundamental basic education, secondary education, and university education. Following the International Society of Ethnobiology Code of Ethics (2006) [40], before we started each survey, the project aims were presented and the consent to use the information was requested from each person. The complete model of the survey is available as an additional file (see Additional file 1).

Questions were categorized into three groups: academic knowledge (1), use and cultural beliefs (2) and interaction and perception (3). The first group refers to questions related to the biology and ecology of the herpetofauna; in the second group the questions were related to the ancestral knowledge of the community about the local herpetofauna and the third group contains questions about perception and human-herpetofauna interactions in daily activities.

### Data analysis

For the obtained data in question groups 1 and 2, a classification of the answers in percentages was made. For the data concerning to question group 3, answers were assigned with a hostility value according to the possible reaction of the individual interviewed in a hypothetical encounter with the herpetofauna. In this way, interaction with amphibians were classified as negative (hostile) behaviors when trying to kill the animal and trying to scare it; neutral behaviors when observing the animal and staying still or moving away; and positive behaviors when transporting the animal to a nearby body of water. Interaction with reptiles were classified in the same way, except for the behavior to remain still or move away, which was considered positive.

Demographic information obtained from the surveys was organized in ordinal categories for each variable. Age was categorized into three groups: (a) young people (14-26 years old), (b) adults (27-51 years old) and elder (52-83 years old). In addition, the data obtained in the urban area of Cumaca were considered as “urban”, whereas the data obtained in the villages of Albania and La Vuelta were considered as “rural”.

In order to know if the positive or negative reactions could be predicted by demographic variables, a Multivariate Ordinal Logistic Regression test (MOLR) was made using the *polr* function of the MASS package in R [41,42]. The response to the interactions with each animal group was defined as the dependent variable and the demographic information as independent variables. The significance of the coefficients obtained from the MOLR was calculated to obtain the *p* values, comparing the *t* values with the standard normal distribution as a Z test [43].

Before making the MOLR test, a covariate correlation test was performed using paired scatter plots in R [44], to assess the possible collinearity of independent variables and eliminate bias in the model. Additionally, the assumption of proportional odds was analyzed using the graphic method recommended by Harrell (2015) [45]. For the validation of the obtained models, the AIC of each model was compared with the null model for each dependent variable (amphibians and reptiles).

In order to compare the samples, proportions differences between the groups were used to assess differences in the answers given the birthplace, education level, and gender; this was done for questions from group 1 (academic knowledge) and 2 (cultural uses and beliefs). The data analysis was done using the software R 3.6.1 [42]. The ubication map of the study area was built with ArcGIS 10.5 [46].

## Results

We surveyed 61 respondents, aged 14-83 with a balanced gender ratio (1:1). Table 1 shows the distribution of the population surveyed in the education level and occupation categories.

**Table 1.** Distribution of the population surveyed into education and occupation categories.

| **Education level** | **Female** | **Male** | **Total** |
| --- | --- | --- | --- |
| High school | 53% | 52% | 52% |
| Primary school | 27% | 32% | 30% |
| College degree | 13% | 6% | 10% |
| Graduate degree | 3% | 3% | 3% |
| Technical studies | 3% | 0% | 2% |
| Illiterate | 0% | 3% | 2% |
| Unanswered | 0% | 3% | 2% |
| **Occupation** |  |  |  |
| Employee, merchant or independent | 27% | 26% | 26% |
| Student | 23% | 29% | 26% |
| Agriculture and land management work | 13% | 32% | 23% |
| Housework and other trades | 37% | 10% | 23% |
| Unemployed | 0% | 3% | 2% |

### Academic knowledge

Regarding amphibians, more than half of the respondents (54.1%) classified frogs and toads within this group; 9.8% classified them as reptiles, whereas 36.1% said they did not know. In addition, they recognized water reserves as a habitat for amphibians, as well as its importance in their life and reproduction cycles. However, only 13% of respondents associated amphibian reproduction with soft egg posture and the presence of larval stages. Most respondents described wet soil, plants with water reserves and crops as a preferred habitat for frogs and toads. Regarding their ecological function, respondents recognized the importance of amphibians as pest controllers and associated their presence with the reduction of mosquitoes’ abundance. Although most of the respondents (70.5%) recognized the existence of poisonous frogs, only 3% stated that it is possible to find them in Colombia.

As for reptiles, 60.7% of respondents classified lizards and snakes within this group, 6.5% classified them as amphibians, whereas 38.8% said they did not know. According to the surveyed population, the main habitats of reptiles are associated with tall trees, crops, grasslands and wet soils. Although the role reptiles play within the ecosystem is unclear, 14.7% of respondents know that they can act as pest controllers.

86.9% of the respondents said that there are venomous snakes in the region. However, only 34.4% of the population surveyed said they had knowledge about the medical treatment associated with snake bites, such as the use of antiophidic serum and the referral to the municipal medical services.

### Use and cultural beliefs

For 63.9% of respondents, amphibians are not part of their cultural or magical-religious beliefs, while 4.9% said they did not know. On the other hand, 31.1% of the population surveyed described cultural beliefs and uses in traditional medicine associated with this group (Table 2). Near 36% of the surveyed population perceive amphibians as dangerous animals, whereas 21.3% describe them as causes of skin diseases such as sores, rash, allergies, inflammations, and even necrosis.

**Table 2.**
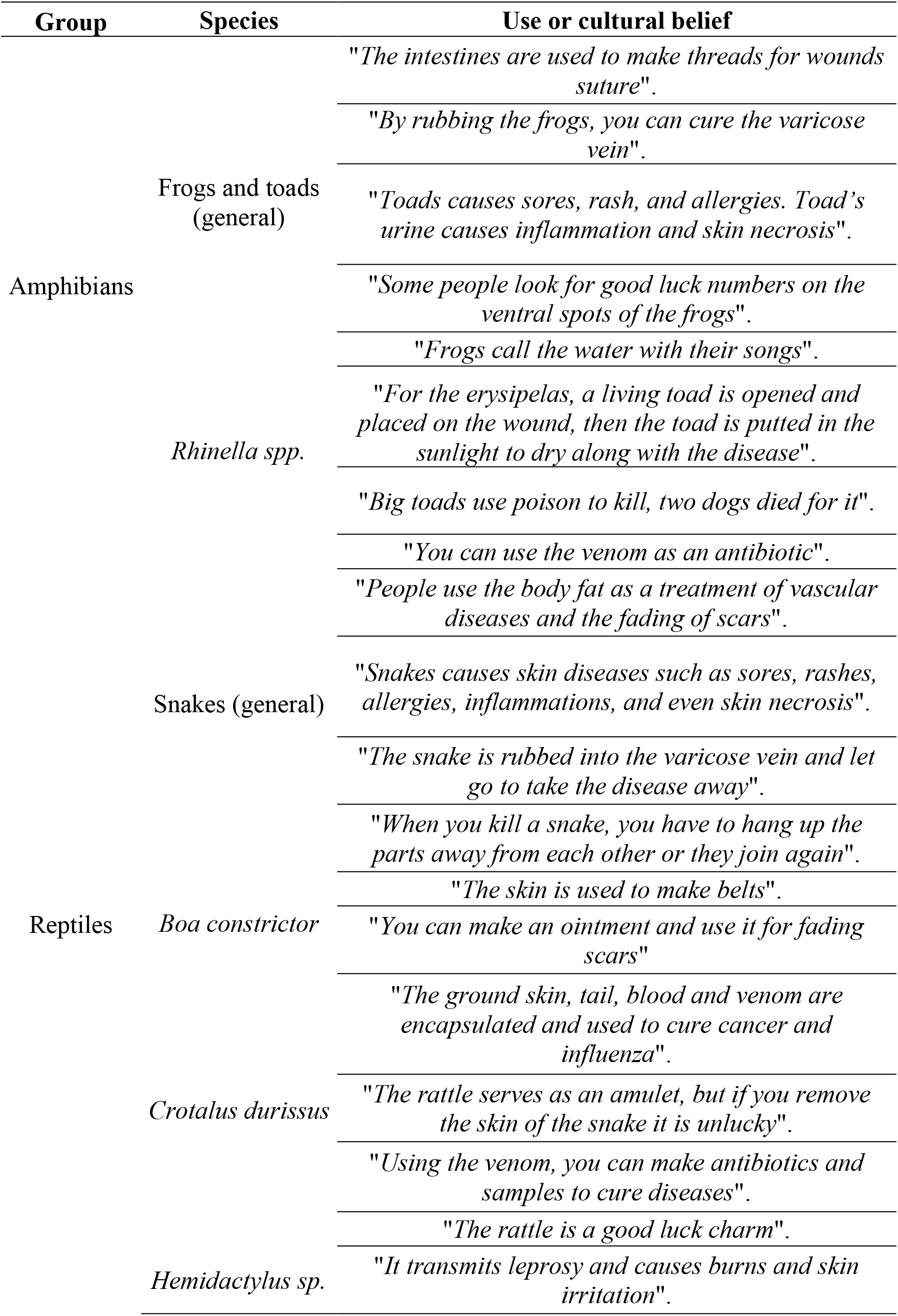
Uses and cultural beliefs of amphibians and reptiles identified in the RFPCQ region.

As for reptiles, 47.5% of respondents did not associate this group with their cultural or magical-religious beliefs, whereas 4.9% said they did not know. 47.5% of respondents described beliefs associated with luck and use in traditional medicine. 88.5% of the population surveyed perceive reptiles as dangerous animals and 27.8% associate them with the transmission of diseases such as leprosy, burns and other skin problems.

### Interaction and perception

The correlation analysis showed that the categories of the occupation presents collinearity with age and level of education, which is why this variable was not considered in the statistical analysis to avoid bias in the model. It was found that the behavior of the data was consistent with the assumption of proportional odds for the variables of origin (rural/urban), gender and age. However, the educational level variable did not fit the assumption, so it was discarded from the model.

The MOLR analysis showed that there is a statistically significant incidence between the variables of origin (MOLR: *t* = 2,378, *p* = 0.017) and gender (MOLR: *t* = −2,322, *p* = 0.020) within the interaction with amphibians, where people living in the urban area are more likely to have a hostile reaction in a possible encounter with an amphibian, compared to people living in the rural area. Also, women tend to have a more negative behavior towards amphibians than to reptiles (Fig. 2).

**Fig. 2.**
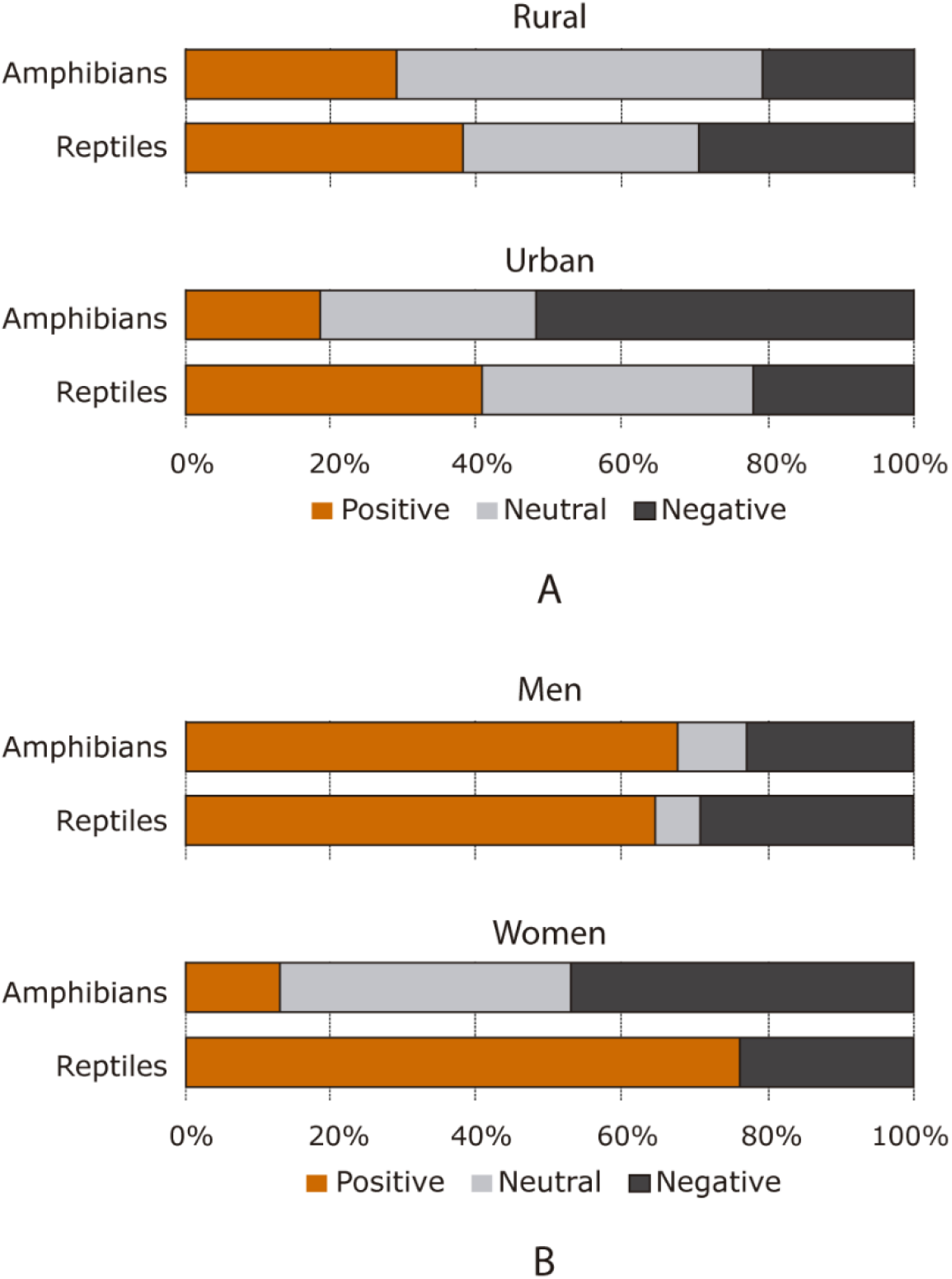
Response tendencies regarding interaction with amphibians and reptiles. It shows the positive, neutral and negative reaction rates in a possible encounter with amphibians and reptiles are shown, according to the origin (**A**) and gender (**B**).

In contrast, there is only a statistically significant incidence for the gender variable (MOLR: *t* = 2,441, *p* = 0.015) in reptile interaction, where men are more likely to react hostile to an encounter with reptiles than women. In addition, there were no neutral behaviors by women in this variable, being the majority positive reactions (76.6%) and the remaining ones negative (23.3%).

The analysis showed that there is no significant relationship between age and the interaction with amphibians and reptiles. In the same way, there was no significant relationship regarding the birthplace and interaction with reptiles (Table 3).

**Table 3.** *p* values obtained in the MOLR analysis regarding the interaction with amphibians and reptiles. Bold: *p* values < 0.05.

| Amphibians |  |  |  |  |
| --- | --- | --- | --- | --- |
| Variable | Factor | Std. Deviation | *t* value | *p* value |
| Origin | Rural ↔ Urban | 0.494 | 2.378 | **0.017** |
| Gender | Male ↔ Female | 0.483 | -2.322 | **0.020** |
| Age categories | Adults ↔ Elder | 0.570 | -0.926 | 0.354 |
|  | Young people ↔ Adults | 0.580 | 0.638 | 0.524 |
| Reptiles |  |  |  |  |
| Variable | Factor | Std. Deviation | *t* value | *p* value |
| Origin | Rural ↔ Urban | 0.520 | 0.693 | 0.488 |
| Gender | Male ↔ Female | 0.499 | 2.441 | **0.015** |
| Age categories | Adults ↔ Elder | 0.587 | 1.573 | 0.116 |
|  | Young people ↔ Adults | 0.602 | -0.396 | 0.692 |

For most people surveyed (55.7%) the abundance of herpetofauna in the RFPCQ has decreased in recent years, especially amphibians. 19.7% of the surveyed population expressed that abundance has increased, whereas 16.4% state that it has remained the same. Finally, 8.2% said they did not perceive a change. Respondents described agricultural activities, chemical fumigation, deforestation, and water pollution as possible current impacts on the amphibian and reptile community in the region. All respondents recognized the importance of generating conservation programs and sustainable usage of the habitat to protect the herpetofauna in the region.

## Discussion

### Academic knowledge

The surveyed population recognizes the existence of amphibians and reptiles in the region, although little is known about general aspects of their biology and the ecological functions. However, the inhabitants properly recognize the potential habitats of these animals, which could help identify priority conservation areas [32,39,47,48].

The local community recognizes the existence of venomous snakes in the region, although it tends to ignore the existence of non-venomous species. This may be due to the association of the body morphology of these animals with risk factors, filing in a general way the serpentiform animals as dangerous; given the evolutionary conditioning of human species [49], which has also been recorded in other primates [50].

Although the population surveyed is aware of the possibility of an ophidic accident, there is a general lack of knowledge about the protocols to follow in case of snake bites, as the use of antiofidic serum is uncommon. This lack of knowledge amplifies the human-snake conflict since the most widespread action to avoid ophidian accidents is killing snakes (even non-venomous), making this one of the main threats to the ophidians population in Colombia [51].

### Use and cultural beliefs

Nearly half of the people surveyed associated reptiles (exclusively snakes) as a part of ancestral culture and traditional medicine, whereas for amphibians this association was lower (31.1%). We propose two reasons as an explanation: 1. As in other regions of Latin America, given the traditional representation of power and the strong ligation of snakes with magical-religious beliefs, the development of a worldview linked snakes to luck and use in traditional medicine is more common, based on the cure through the power and the spirit that snakes represent [52]. 2. Since amphibians are mainly nocturnal, a meeting with people in their daily activities is less common. This has resulted in a less strong interaction and a minor influence on the worldview of the inhabitants of the region. Although Panche indigenous communities made drawings of amphibians and associated them with water [5], it seems that these traditions were not maintained over time in the current inhabitants.

A third of the population surveyed perceives amphibians as dangerous and disease-causing animals. The main belief was that skin wounds are caused by urine or secretions of the parotid glands of the toads (*Rhinella* spp.). Although information on the effect of bufotoxins on human health is scarce [53,54], there are extensive records of intoxication in dogs [55–58], and wild vertebrates [59–61]. This is a problem especially widespread in countries in which this species has been introduced, sometimes causing the death of the animals involved [56,58]. However, it is more common to find that toads are avoided due to their physical appearance, which, added to the lack of knowledge about their biology, results in the association of these animals with areas of debris and dirt.

On the contrary, most of the population surveyed perceive reptiles as dangerous animals (88.5%), based mainly on the fear of an ophidian accident. Although 47.5% of the reported cases of poisoning in Colombia correspond to ophidian accidents [62], between 2014 and 2016 only 185 cases of snakebite were reported in the department of Cundinamarca (61 cases on average per year; [63]), being one of the departments with the lower reported cases in Colombia (< 2%). This can indicate that the response of fear towards snakes is more linked to the cultural imprint than to the interaction and previous experiences of the inhabitants with the snakes in the area.

Despite this, there are several uses of amphibians and reptiles in traditional medicine, such as the use of dissected toads for the cure of varicose veins, shingles (herpetic skin lesion), erysipelas and other dermal bacterial infections; and the use of snakes for the cure of some diseases such as cancer and heart ailments (Table 2). Previous studies have also reported these uses in other Latin American countries [64,65] and other regions of the world [66,67]. In addition, in recent decades there has been increasing evidence on the possibility of pharmacological synthesis based on chemical components obtained from secretions, tissue and venom extracted from amphibians and reptiles [68–71], which shows the correspondence between the use of species in the traditional culture, as well as in current and future applications in medicine.

### Interaction and perception

Regarding the response of the rural population towards the interaction with amphibians is dominated by neutral and positive behaviors, with a lower incidence of negative behaviors, which contrasts with the response to the same interaction in the urban population (Fig. 2)

For the rural population amphibians are a fundamental part of the nocturnal sound ecosystem, which could have a positive impact on the perception towards these animals. Being less frequent the occurrence of amphibians in the urban area, it is likely that people develop a negative perception due to the lack of contact. On the contrary, the trend response to interaction with reptiles between the rural and urban population is similar, which could indicate that there is an imprint of information on reptiles that remains more stable between the community and birthplaces.

Despite people’s cultural/natural fear towards snakes [49,72], in most cases, the positive reactions in reptiles interaction were higher regarding the interaction with amphibians. However, this may be due to the difference in the classification of positive and neutral reactions, where the behavior of standing still or moving away is considered neutral for interaction with amphibians, whereas it is positive for interaction with reptiles. The above is based on the idea that possible positive behaviors towards reptiles (specifically snakes) do not consider transporting the animal to a safe place, due to its potential danger [72–74].

Men described mostly positive behaviors in response to interaction with amphibians and reptiles, tending to have few neutral behaviors in both groups. This indicates that, although snakes are perceived as dangerous animals, this usually does not lead to defensive behavior that implies the threat of species in the study region, a phenomenon previously reported in other parts of Latin America [72,74]. In contrast, women in the surveyed population tend to have more negative behaviors in the interaction with amphibians with respect to reptiles. This may be related to the possible danger of snakes, since taking a negative reaction and attacking the animal implies a health risk, which does not occur with amphibians and could make them more susceptible to be attacked.

Although the data provided by this study represent a preliminary approach to the understanding of current human-herpetofauna interactions in the RFPCQ, the implementation of a more direct methodological design is necessary, in order to test the reactions of people in a controlled environment. These studies are vital as a basis for planning conservation strategies in the region since the interaction of humans with nature directly influences the survival dynamics of current animal populations [75,76].

### Final considerations

When local communities are empowered with basic knowledge about the species that occur in their region and become active agents of participation for their conservation, management, and preservation, plans are effective [47,48,77]. Local population recognize the importance of preserving and conserving nature, although they are unaware of the fundamental reasons why this type of action should take place. In this way, processes such as deforestation, degradation, and habitat loss caused mainly by human activities such as improper use of land, overexploitation of natural resources and the exclusivity of land use for agriculture and cattle ranching need to be reassessed [76,78].

Due to the current way in which science communicates, where the vast majority of scientific papers are not very informative to local communities, it is usual to find out that local populations have a limited scope of knowledge, which does not allow its use in decision making [79]. Although biology is one of the disciplines that use the most technical terms, efforts to ensure assertive communication must be prioritized, which allow reducing sectorization and making information available to people most directly related to the natural environment [80].

## Conclusion

Although there was a general lack of knowledge on the part of the inhabitants of the RFPCQ about the biological and ecological aspects of herpetofauna, the population recognized the basic information about the habitats of these animals within the area of the reserve, which could help identify possible priority conservation areas.

Most of the population do not associate the herpetofauna with cultural or magical-religious traditions. However, there is a wide variety of uses of amphibians and reptiles in traditional medicine, which are consistent with the information obtained in other regions of Latin America and the world.

In general, the perception of amphibians and reptiles varied according to the origin and gender of the people surveyed, tending to have a more positive perception about reptiles that of amphibians in most cases.

Greater efforts should be made in the transmission and dissemination of knowledge about the ecological functions of herpetofauna, which will contribute to the success of future conservation plans in the area with the local community participation.

## Acknowledgments

We would like to thank Castillo family, Quininí farm and to the Asociación de Protectores de los Recursos Naturales y del Ambiente de Tibacuy (APRENAT) association for their hospitality, logistic help and compromise with the academic activities at the RFPCQ. Thanks go to Felipe Escobar for his help in the construction of the statistical models and revision of the manuscript and to Andrés Olaya for his help in preparing the RFPCQ location map. Also, we want to thank to Uber Rozo, Miller Castañeda, Johana Muñoz, Miguel Méndez, Jherandyne Castillo, Juliana Poveda and Sebastián Pérez for their help in the fieldwork and participation in different stages of the project. To the Grupo estudiantil de Herpetología de la Universidad Nacional de Colombia (HERPETOS UN) and its members for helping us with fieldwork and the tabulation of the surveys. We thank the Project Management Program (PGP – Bienestar Universitario) and Facultad de Ciencias of Universidad Nacional de Colombia, Sede Bogotá, who financed the field stage and information gathering. Also, thanks to Dr. Martha Calderón and Dr. Adriana Jerez for the revision and invaluable comments of the manuscript.

## Abbreviations

MOLR: Multivariate Ordinal Logistic Regression
RFPCQ: “Reserva Forestal Protectora Cerro Quininí” forest reserve
Sp (p): Species

## Declarations

### Authors’ contributions

All authors worked equally on this manuscript.

### Availability of data and materials

Complete survey model can be found as Additional file 1. Survey format conducted on the inhabitants in the study area.docx.

Data collected from surveys are available in Additional file 2. Surveys data.xlsx.

### Ethics approval and consent to participate

The study conforms to the code of ethics of the International Society of Ethnobiology [40]. Prior informed consent was obtained from all participants before interviews.

### Consent for publication

Not applicable.

### Authors’ information

J. C Ríos-Orjuela is a biologist from the National University of Colombia and an MSc student in systematics and biodiversity at the University of São Paulo. His interests include herpetology and ornithology and his research works include aspects of functional morphology in reptiles, evolution of neotropical birds and citizen science education.

N. Falcón-Espitia is a biologist from the National University of Colombia. His interests are based on herpetology and his research works include aspects of morphology in reptiles and ecology.

A. Arias-Escobar is a biologist from the National University of Colombia. Her interests include ecology, morphology, and conservation of amphibians and reptiles and citizen science education.

M. J. Espejo-Uribe is a biologist from the National University of Colombia. Her general interests are ecology and conservation genetics, taking as principal research groups amphibians and birds.

C. T. Chamorro-Vargas is a bachelor’s student in biology from the National University of Colombia. Her interests are herpetology and ecology, and her research works include aspects of life history of reptiles and ecoacustics of amphibians.

